## Supplementary Material for "Neonatal-derived IL-17 producing dermal γδ T cells are required to prevent spontaneous atopic dermatitis"

### Supplemental Figure Legends

**Supplemental Figure 1**. Enhanced effector function of ILC and Tγδ17 cells in mice with genetic propensity for skin inflammation and cytokine milieu that propagate altered innate lymphocyte hyperactivity.

A) Gating scheme for FACS analysis of ILC in the skin.

B) Analysis of IL-17A and IL-13 production by ILC from 3mo mice as identified in Panel A after PdBU/ionomycin stimulation.

C) Intranuclear GATA3 staining in skin ILC. Non-T cells and T cells are presented as internal references for the high level of GATA3 expressed by ILC

D) Increased skin γδ T cells, including Tγδ17 cells ($TCR\delta^{int}$), in 4-6 week old *Rora*$^{-/-}$ mice. Skin cells were stimulated with PdBU/ionomycin and IL-17 production assessed in CD3+ T cells. One of 4 mice analyzed shown. Note limited IL-17 production at this age from αβ T cells and no IL-17 production from $TCR\delta^{hi}$ $V\gamma3^{+}$ dendritic epidermal T cells (DETCs).

E) RNA was prepared from total muzzle skin (6 mo mice) and analyzed by RT-qPCR for cytokine and chemokine gene expression. Only those significantly altered in expression (except *Il1a* as an example of unchanged) are shown. LMC n=4; *Sox13*$^{-/-}$ mice, n=5. ***, p<.001; *, p<.05 by Student's *t*-test.

**Supplemental Figure 2.** Characterization of skin MAITs, iNKT cells, regulatory T cells (Tregs), B cell subsets and enhanced activation of melanocyte antigen-specific T cells in *Sox13*$^{-/-}$ mice.

A) Control MAIT tetramer (MR1/6-FP) staining (left two panels), and MR1/5-OP-RU (5-(2-oxopropylideneamino)-6-Dribitylaminouracil, a riboflavin biosynthetic product recognized by MAITs) staining of skin among $CD4^{+}$ and $CD8\beta^{+}$ T cells reveal minimum coreceptor expressing MAITs in the skin of 5-6mo mice.

B) Analysis of MAIT cells in skin dLN from mice as above, including coreceptor and CCR6 expression, a hallmark of type 3 cytokine producing T cells.

C) Analysis of iNKT cells in the skin and dLN of mice in panel A, using CD1d/PBS57 tetramer that stains the canonical Vα14+ iNKT cells.

D) Intranuclear staining for FoxP3 to identify Tregs in the ear skin of 5-6 mo LMC (top) and *Sox13*$^{-/-}$ (bottom) mice. No changes in frequency of FOXP3+ cells were observed in the muzzle skin.

E) Representative FACS analysis of GC B cells (GL7+ CD95+) in dLN of indicated 5 mo mice.

F) Representative FACS analysis of CD138+ B220- plasma cells in dLN of indicated 5 mo mice.

G) Representative FACS analysis of PD-1hi Bcl6+ Tfh in dLN of indicated 5 mo mice.

H) Pre-melanocyte antigen 1-specific PMEL CD8+ Tg T cells were labeled with CellTrace Violet and then adoptively transferred into 6 mo LMC or *Sox13*$^{-/-}$ mice. Four days later, spleen and skin dLN were harvested for analysis of proliferation by transferred PMEL T cells (identified by Thy1.1 allomarker). Data from one of two experiments shown, n=4/group.

I) Summary data of PMEL adoptive transfer proliferation performed in Panel H. **, p<.01; *, p<.05 by ANOVA.

**Supplemental Figure 3.** Skin microbiome of Abx *Sox13*$^{-/-}$ mice and independence of dermal nTγδ17 from CB for their development and maintenance.

A) Muzzle skin microbiome analyses of Abx *Sox13*$^{-/-}$ mice at 3 and 6 mo show near complete depletion of *Corynebaceria mastitidis* and decreased *Staphylocccus*, compared to the age-dependent blooming of these species in *Sox13*$^{-/-}$ mice shown in Fig. 5, and correlating with the prevention of AD in Abx *Sox13*$^{-/-}$ mice.

B) Loss of fTγδ17 cells (CD3$^{hi,}$ among TCRδ$^{+}$ cells) in 6 wk old germ free (GF) B6 mice in dLN and skin, relative to those housed in the SFP condition, as reported previously. However, nTγδ17 cells are maintained and fully capable of IL-17 production in age-matched GF mice. Representative of 4-5 mice/group from two independent experiments.

C) Abx *Il17a*-Egfp mice show that in vivo *Il17a* transcription in nTγδ17 cells occurs independent of the Abx sensitive CB in panel A. Mice were treated with Abx for 4 weeks, and then ex vivo dermal γδ cells in the skin (TCRδ$^{int}$) were analyzed for expression of Vγ chain (left panel, Vγ2$^{neg}$ dermal γδ T cells are Vγ4$^{+}$) and CCR6 and EGFP (right two panels) expression in dermal Vγ2$^{+}$ nTγδ17 and Vγ2$^{-}$ (Vγ4$^{+}$) fTγδ17 cells.

D) Summary of data from experiments depicted in Panel C show the loss of fTγδ17 cells and compensatory increase in nTγδ17 cells in 1 month Abx mice (Top) and the preserved capacity to express *Il17a* in nTγδ17, but not in residual fTγδ17, cells. n=3/group. ***, p<.001 by ANOVA.

**Supplemental Figure 4.** Expansion of CD4$^{+}$Vβ4$^{+}$ T cells in the skin of *Sox13*$^{-/-}$ mice.

A) Representative multiplex analysis of TCR Vβ usage in skin T cells of 13 Vβ for which commercial Abs are available. In this illustration, in Panel A αVβ4 and αVβ9 are both FITC conjugates; αVβ6 and Vβ12 are PE conjugates; αVβ2, αVβ4, and αVβ6 are biotin conjugates. This combinatorial staining strategy allows for deconvolution of which TCR Vβ is expressed by a given population via the gating shown. Inset red numbers in parentheses indicate the specific Vβ identified by the gated population.

B) Heatmap of TCR Vβ expression by T cells in the skin of LMC and *Sox13*$^{-/-}$ mice. n=9, pooled from 3 independent experiments using 6mo mice. From this analysis, only Vβ4 in CD4$^{+}$ cells was consistently and significantly altered across multiple experiments. N.R., non-reactive to Abs in panel. Color coding at the bottom indicates % positive among αβ T cell subsets. ***, p<.001 by ANOVA

C) Intranuclear FoxP3 staining in skin CD4$^{+}$Vβ4$^{+}$ T cells from 5-6mo mice. Compared to Supp Fig. 2D, data indicate that CD4$^{+}$Vβ4$^{+}$ T cells are underrepresented in the skin Treg pool.

D) Intranuclear expression of Ki-67 in skin CD4$^{+}$Vβ4$^{+}$ T cells from 5-6mo mice , indicating their enhanced proliferation in *Sox13*$^{-/-}$ mice. n=3 from 1 of 2 similar experiments. *, p<.05 by Student's *t*-test.

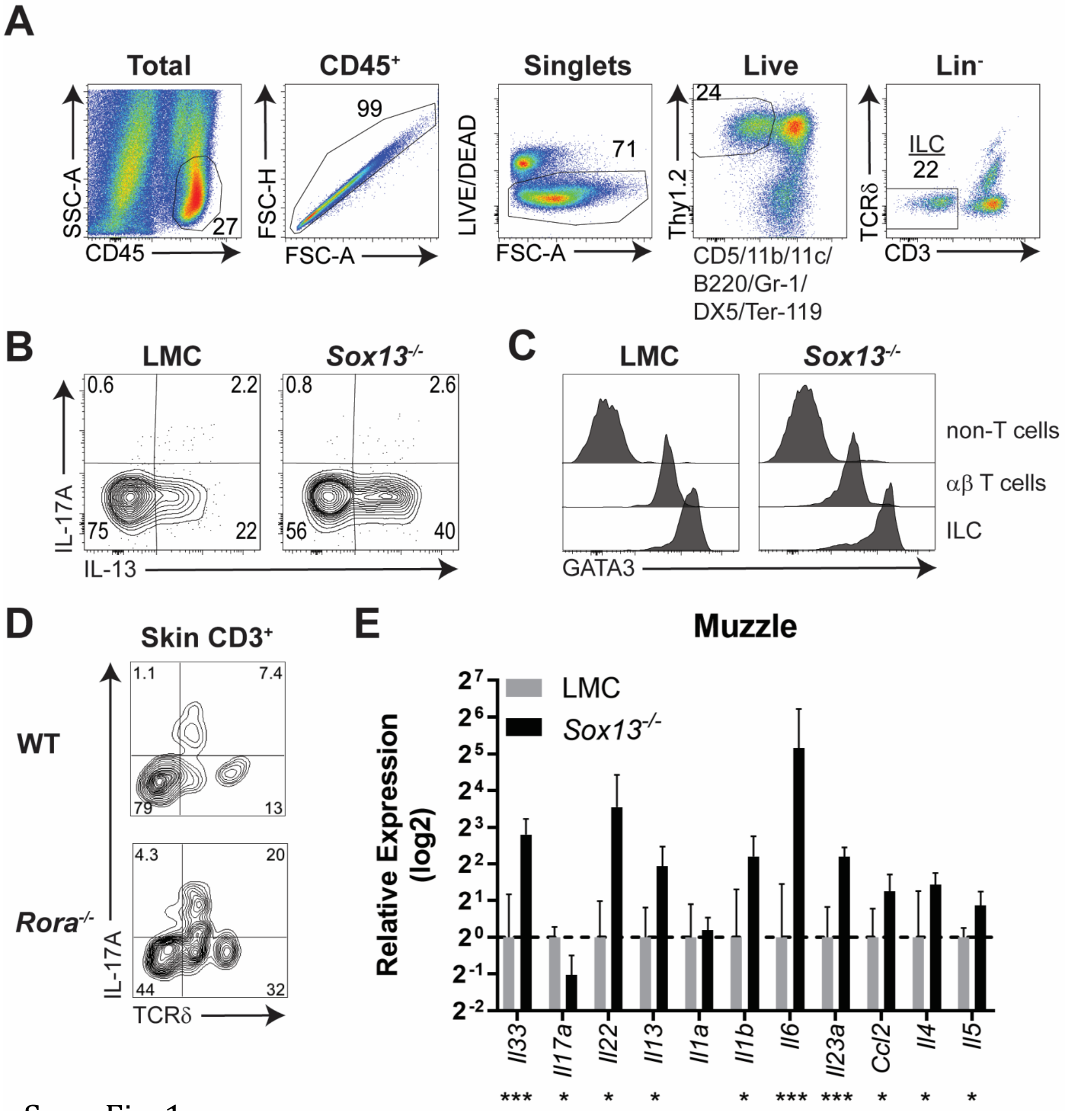
A
Total
CD45+
Singlets
Live
Lin-
SSC-A
CD45
27
FSC-H
FSC-A
99
LIVE/DEAD
FSC-A
71
Thy1.2
CD5/11b/11c/
B220/Gr-1/
DX5/Ter-119
24
TCRδ
CD3
ILC
22
B
LMC
Sox13-/-
IL-17A
IL-13
0.6
2.2
75
22
0.8
2.6
56
40
C
LMC
Sox13-/-
non-T cells
αβ T cells
ILC
GATA3
D
Skin CD3+
WT
Rora-/-
IL-17A
TCRδ
1.1
7.4
79
13
4.3
20
44
32
E
Muzzle
LMC
Sox13-/-
Relative Expression
(log2)
Il33
Il17a
Il22
Il13
Il1a
Il1b
Il6
Il23a
Ccl2
Il4
Il5
*** * * * * *** *** * * *

Supp Fig. 1

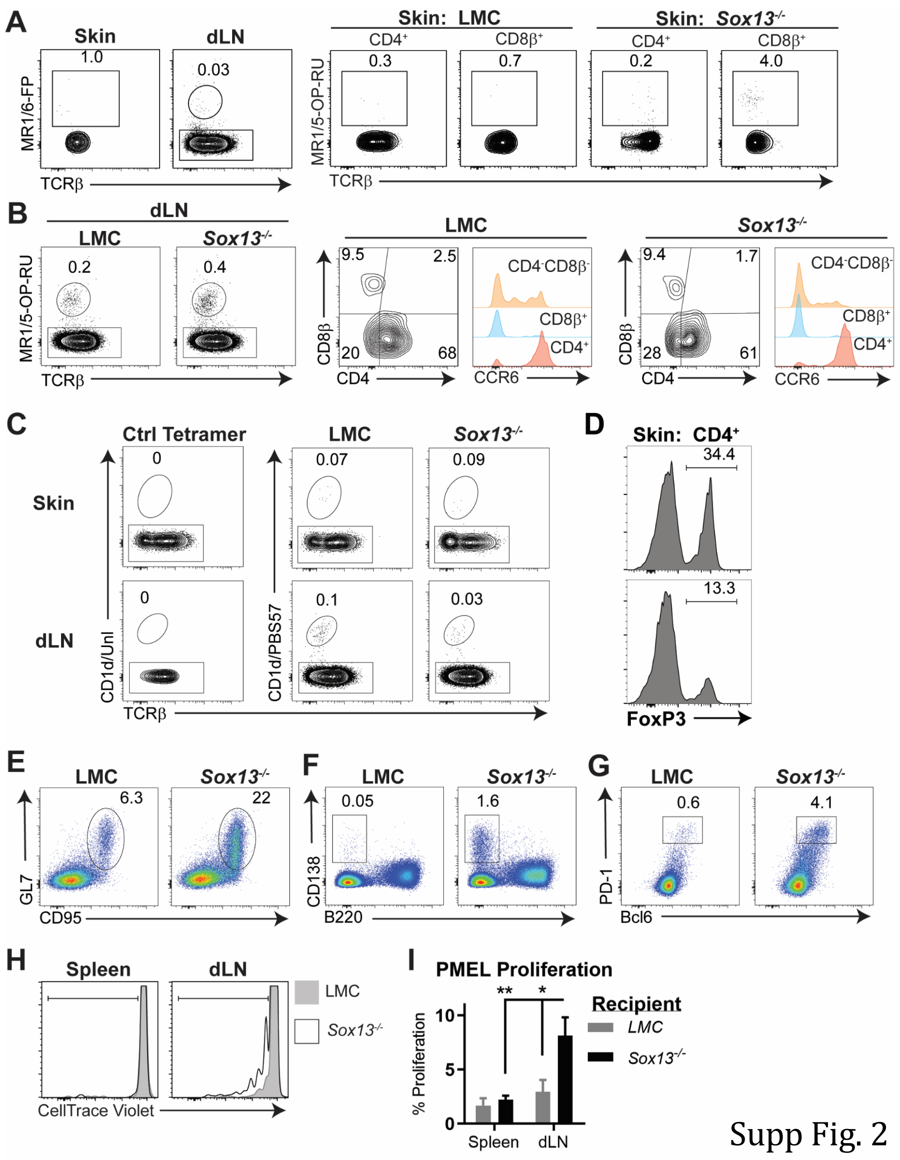
A
Skin
dLN
1.0
0.03
MR1/6-FP
TCRβ
Skin: LMC
Skin: Sox13-/-
CD4+
CD8β+
0.3
0.7
0.2
4.0
MR1/5-OP-RU
B
dLN
LMC
Sox13-/-
0.2
0.4
9.5
2.5
20
68
9.4
1.7
28
61
CD8β
CD4
CCR6
CD4-CD8β-
C
Ctrl Tetramer
Skin
dLN
0
0.07
0.09
0.1
0.03
CD1d/Unl
CD1d/PBS57
D
Skin: CD4+
34.4
13.3
FoxP3
E
6.3
22
GL7
CD95
F
0.05
1.6
CD138
B220
G
0.6
4.1
PD-1
Bcl6
H
Spleen
CellTrace Violet
I
PMEL Proliferation
Recipient
% Proliferation
**
*
10
5
0

Supp Fig. 2

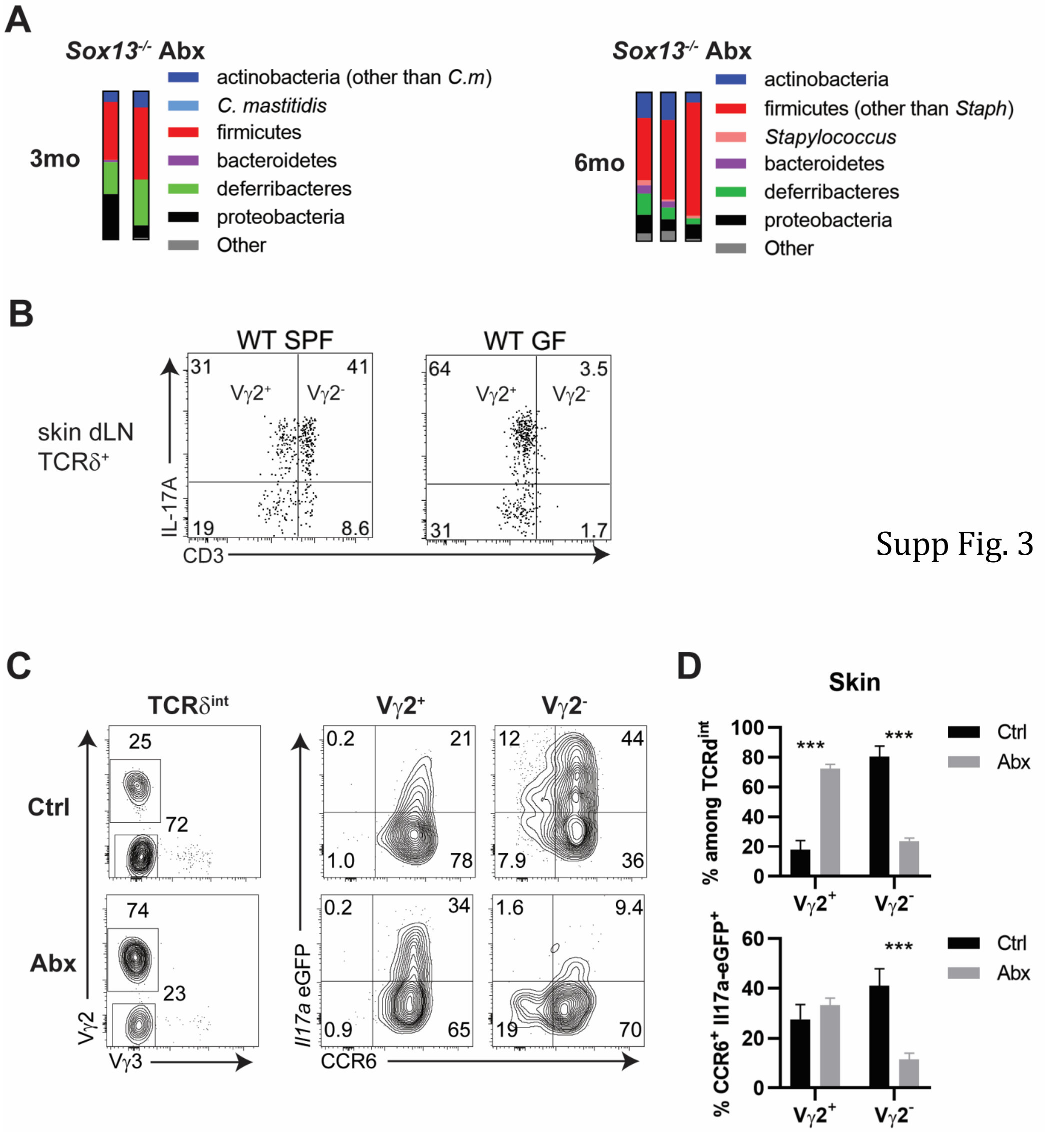

A
Sox13-/- Abx
3mo
actinobacteria (other than C.m)
C. mastitidis
firmicutes
bacteroidetes
deferribacteres
proteobacteria
Other
Sox13-/- Abx
6mo
actinobacteria
firmicutes (other than Staph)
Stapylococcus
bacteroidetes
deferribacteres
proteobacteria
Other
B
WT SPF
WT GF
skin dLN
TCRδ+
IL-17A
CD3
Vγ2+
Vγ2-
31
41
19
8.6
64
3.5
31
1.7
C
TCRδint
Ctrl
Abx
25
72
74
23
Vγ2
Vγ3
Vγ2+
Vγ2-
0.2
21
1.0
78
12
44
7.9
36
0.2
34
0.9
65
1.6
9.4
19
70
Il17a eGFP
CCR6
D
Skin
% among TCRdint
% CCR6+ Il17a-eGFP+
Ctrl
Abx
***
Vγ2+
Vγ2-
100
80
60
40
20
0


Supp Fig. 3

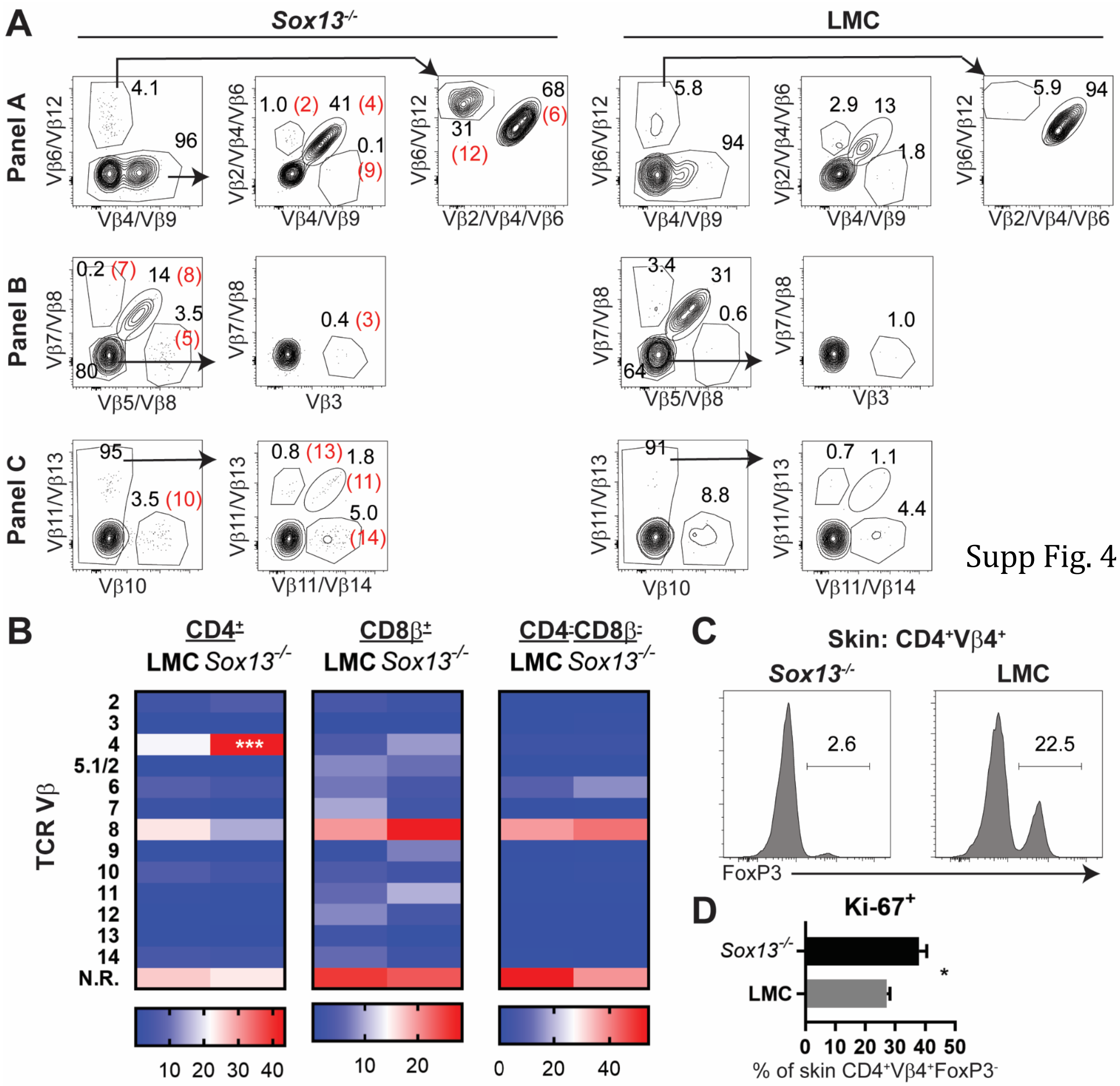
A
Sox13-/-
LMC
Panel A
Panel B
Panel C
B
CD4+
CD8β+
CD4-CD8β-
LMC Sox13-/-
TCR Vβ
C
Skin: CD4+Vβ4+
FoxP3
D
Ki-67+
% of skin CD4+Vβ4+FoxP3-

Supp Fig. 4

Supplemental Table 1. PCR Primers used in this study

| Sequence | F/R | Description |
|---|---|---|
| CCTGGACTCTCCACCGCAA | F | Il17a |
| TTCCCTCCGCATTGACACAG | R | Il17a |
| TTTCCTGTCTGTATTGAGAAACCT | F | Il33 |
| TATTTTGCAAGGCGGGACCA | R | Il33 |
| CGCTTGAGTCGGCAAAGAAAT | F | Il1a |
| TGGCAGAACTGTAGTCTTCGT | R | Il1a |
| GCCACCTTTTGACAGTGATGAG | F | Il1b |
| GACAGCCCAGGTCAAAGGTT | R | Il1b |
| TCCTCTCTGCAAGAGACTTCC | F | Il6 |
| TTGTGAAGTAGGGAAGGCCG | R | Il6 |
| AGCTGTAGTTTTTGTCACCAAGC | F | Ccl2 |
| GTGCTGAAGACCTTAGGGCA | R | Ccl2 |
| TCACAGCAACGAAGAACACCA | F | Il4 |
| CAGGCATCGAAAAGCCCGAA | R | Il4 |
| CAAGCAATGAGACGATGAGGC | F | Il5 |
| GCATTTCCACAGTACCCCCA | R | Il5 |
| CACTACGGTCTCCAGCCTCC | F | Il13 |
| CCAGGGATGGTCTCTCCTCA | R | Il13 |
| CACCAGCGGGACATATGAATCT | F | Il23a |
| CACTGGATACGGGGCACATT | R | Il23a |
| AGAGTTTGATCCTGGCTCAG | F | 16S V1 Universal Primer 27F |
| ATTACCGCGGCTGCTGG | R | 16S V3 Universal Primer 534R |
| AAGCCTGATGACTCGGCCACA | F | Vb4 TCR deep seq |
| CTTGGGTGGAGTCACATTTCTCAGATCCTC | R | Cbeta TCR deep seq |
| AACTGTACTTATTCAACCACA | F | Va4 TCR deep seq |
| CTGTGAACTGTTCCTATGAAACC | F | Va4 TCR deep seq |
| TAAACTGTACTTATTCAACCACA | F | Va4 TCR deep seq |
| CCTGATAATAAATTGCACGTATTCA | F | Va4 TCR deep seq |
| GGTACACAGCAGGTTCTGGGTTCTGGATG | R | Calpha TCR deep seq |
